# Root water gates and not changes in root structure provide new insights into plant physiological responses and adaptations to drought, flooding and salinity

**DOI:** 10.1101/2020.10.27.357251

**Authors:** Jean-Christophe Domec, John S. King, Mary J. Carmichael, Anna Treado Overby, Remi Wortemann R, William K. Smith, Guofang Miao, Asko Noormets, Daniel M. Johnson

## Abstract

The influence of aquaporin (AQP) activity on plant water movement remains unclear, especially in plants subject to unfavorable conditions. We applied a multitiered approach at a range of plant scales to (i) characterize the resistances controlling water transport under drought, flooding and flooding plus salinity conditions; (ii) quantify the respective effects of AQP activity and xylem structure on root (K_root_), stem (K_stem_) and leaf (K_leaf_) conductances, and (iii) evaluate the impact of AQP-regulated transport capacity on gas exchange. We found that drought, flooding and flooding-salinity reduced K_root_ and root AQP activity in *Pinus taeda*, whereas K_root_ of the flood-tolerant *Taxodium distichum* did not decline under flooding. The extent of the AQP-control of transport efficiency varied among organs and species, ranging from 35%-55% in K_root_ to 10%-30% in K_stem_ and K_leaf_. In response to treatments, AQP-mediated inhibition of K_root_ rather than changes in xylem acclimation controlled the fluctuations in K_root_. The reduction in stomatal conductance and its sensitivity to vapor pressure deficit were direct responses to decreased whole-plant conductance triggered by lower K_root_ and larger resistance belowground. Our results provide new mechanistic and functional insights on plant hydraulics that are essential to quantifying the influences of future stress on ecosystem function.

## Introduction

There is scientific consensus that the Earth’s climate is changing at a geologically unprecedented rate and that human activities are a contributing factor, indicated by the National Academy of Sciences survey on climate change (NAS, 2020), and bolstered by a recent IPCC report (Oppenheimer et al., 2019). Due to a combination of seawater thermal expansion and melting of glaciers and polar ice sheets, global sea level rose 0.17 m over the 20^th^ century and is projected to rise by at least 0.35 m by 2100 (Peltier, 2002). Coastal forests are among the world’s most biologically diverse and productive ecosystems, but unfortunately are also the most vulnerable to sea level rise (SLR; Kirwan and Gedan, 2019). In addition to increased flooding, SLR is globally expected to foster high salinities into tributary freshwater areas of the coastal zones (Bhattachan et al., 2018). At the same time of being subject to increased salinity, those threatened ecosystems undergo periodic droughts exposing coastal forests to low soil water availability (DeSantis et al., 2007). Understanding forest responses to SLR therefore requires the determination of physiological response mechanisms to drought, flooding and flooding plus salinity.

Scientists have a broad-scale understanding of plant adjustment and tolerance to flooding and salinity along environmental gradients and the bulk of recent work in plants has been crucial in distinguishing adaptive plant strategies (Kirwan and Gedan, 2019). One of the most characteristic traits of wetland plants is aerenchyma, a specialized tissue made of intercellular gas-filled spaces that improves the storage and diffusion or oxygen. Overall, the physiological responses of plants to salt stress and flooding are similar in many ways (Allen et al., 1996; Munns, 2002), but the mechanisms by which plants deal with these stressors differ across species (Munns and Tester, 2008). The main consensus is that the primary responses of plants to flooding and salt is inhibition of root hydraulic conductance (Loustau et al., 1995; Rodríguez-Gamir et al., 2012).

In turn, this reduction in water uptake capacity reduces photosynthesis and growth due to the closure of stomata (McLeod et al., 1996; Munns and Tester, 2008). However, there are no studies that have focused on variation in hydraulic traits in contrasting species in terms of adaptive strategies and root physiological responses to full inundation, limiting our understanding of how they are linked to leaf- and whole-plant-level water transport, which limits our ability to predict forest ecophysiological response to SLR and climate change.

Water flow in the soil-plant-atmosphere continuum (SPAC) is determined by the hydraulic conductance of soil and plant tissues, which characterizes the structural capacity of the whole plant to move water (Tyree and Zimmermann, 2002). Hydraulic conductance (K_plant_) is an important factor predicting gas exchange, transpiration, plant water status, growth rate and resistance to environmental stresses (Sperry, 2003; Addington et al., 2004; Brodribb and Holbrook, 2003; McCulloh et al., 2019). The partitioning of K_plant_ along the water transport path is very variable, not only among species, but also diurnally and among plant organs (Ye and Steudle, 2006; Johnson et al., 2016). Approximately 50-60% of the whole-plant hydraulic resistances (1/K_plant_) are located in the root system, which shows the outstanding importance of this organ within the flow path (see review by Tyree and Zimmermann 2002). Peripheral organs such as leaves and roots have been proposed as possible replaceable hydraulic fuses of the SPAC during stress, uncoupling stems hydraulically from transpiring surfaces and soil (Hacke et al., 2000; Sperry, 2003; Domec et al., 2009 Johnson et al., 2016). Quantifying the relative contribution of K_root_ to K_plant_ and how it varies under drought, flooding, and flooding plus salinity is thus essential for understanding how these stressors influence photosynthesis and stomatal conductance (g_s_) and their sensitivity to climatic variables. In addition, mechanistic depiction of variation in K_plant_ and its impact on *g*_s_, and the sensitivity of g_s_ to prominent environmental drivers, requires isolating the main resistances to plant water flow, and their dynamics in response to abiotic stress factors, which has rarely been done.

Most research on abiotic stresses has focused on aboveground organs and neglected physiological responses of the roots, especially in woody plants. This is surprising because important processes of plant tolerance are located in the roots and also because roots are the first organs to be affected by water stress, flooding and salinity (Krauss et al., 1999). In radial and axial roots axes, resistance to water flow depends on root anatomy (Knifer and Fricke, 2011), whereas in the radial component it is also a function of protein water channels, or aquaporins (AQP), that regulate the resistance of the transcellular pathway (Chaumont et al., 2005; Gambetta et al., 2017). Aquaporins are imbedded in the plasma and vacuolar membranes of most root cell types and form pores that are highly selective for water (Tornroth-Horsefield et al., 2006). In crop plants AQP chemical inhibitors (i.e. mercuric chloride or hydroxyl radicals) demonstrated that AQP down-regulation is the principal cause of alterations of the radial pathway, resulting in a decrease in K_root_ (Ehlert et al., 2009; Knifer and Fricke, 2011; Maurel and Nacry 2020). Despite a recent flurry of studies, compared to reference plants used in molecular studies such as corn, tobacco and *Arabidopsis* (Siefritz et al., 2002; Lopez et al., 2003; Bramley et al., 2009; Sade et al., 2010; Tan and Zwiazek, 2019), the importance of species differences in AQP regulation in woody plants and its effect within the SPAC is still poorly understood (McElrone et al., 2007; Gambetta et al., 2013; Johnson et al., 2014; Rodriguez-Gamir et al., 2019).

Better information on physiological functioning of forest species in stressed environments is needed to develop adaptive management strategies that will help protect threatened coastal ecosystems (Carmichael and Smith, 2016). To fully understand the impacts of SLR on plant adaptation, the influence of abiotic stresses on root hydraulics must be evaluated with respect to the entire capacity of the plant to move water. This is especially relevant for seedlings, which have low physiological capacity to tolerate many stressors (Niinemets, 2010), impacting species persistence under changing conditions (Megonigal and Day, 1992; Brodersen et al., 2019). In that framework, our first objective was to characterize the vascular conductances that control water movement through the plant system under drought, flooding, and flooding plus salinity stresses. Our second objective was to quantify the effects of AQP activity on plant organs and partition the antagonistic effects of AQP and xylem structure on conductances. Our third objective was to evaluate a hypothesized correlation between leaf-level gas exchange and AQP regulation of water transport under varying environmental conditions. Using contrasting species, we tested the hypotheses that decrease in hydraulic conductance between treatments 1) is controlled by AQP activity rather than by a change in root xylem structure with greater declines in stress-intolerant plants, and in roots than in stems and leaves; and 2) is optimized in plants experiencing lower AQP inhibition, such that K_root_ exerts greater control on K_plant_, which in turn affect g_s_ and carbon assimilation when environmentally stressed.

## Material and Methods

### Plant material and greenhouse experiments

We used 50 one-year-old *Taxodium distichum* L. and *Pinus taeda* L. half-sib seedlings supplied by ArborGen Inc. (Ridgeville, SC, USA). At the beginning of the spring season (late March), the seedlings were repotted in 19 liters commercial plant pots filled with a Fafard-4P soil mixture composed of sphagnum peat moss (50%), bark (25%), vermiculite (15%) and perlite (Fafard Inc., Agawam, MA, USA). This mixture was representative of the soil texture and organic matter content of soils found in coastal forested wetlands. Potted plants were maintained in a greenhouse with a 16-h photoperiod where daytime mean temperature and relative humidity were kept at 23±3°C and 55±6 %, respectively. Before the treatments were applied, all 50 plants were watered three times a week. Eight weeks after the beginning of the experiment 36 plants were randomly separated into four groups (control, droughted, flooded, flooded plus salt) and were surrounded and buffered by the 14 plants that were not used for the measurements. These treatments were intended to represent stresses related to SLR and periodic droughts exposing coastal forests, and thus the soil salinity treatment with no flooding was not studied. Those single-factor experiments were conducted simultaneously and applied for 35 days (Rodriguez-Gamir et al., 2019). Control plants were irrigated with 2 liters of water twice per week, which was enough to saturate the substrate. For the drought treatment, plants were never irrigated from the start until the end of the experiment (Rodriguez-Gamir et al., 2019). Flooding and flooding plus salinity was imposed by submerging the seedlings to the root-collar (3 cm above the surface) without draining the pots (Pezeshki, 1992). The salinity treatment (at a concentration of 4 g 1^−1^, or 4 parts per thousand) was prepared using a commercial seawater mixture (24 g 1^−1^ NaCl; 11 g 1^−1^ MgCl_2_-6H_2_O; 4 g 1^−1^ Na_2_SO_4_; 2 g 1^−1^ CaCl_2_-6H_2_O; 0.7 g 1^−1^ KCl).

### Hydraulic conductance of root, shoot and whole plant

Five weeks after the treatments were applied, root (K_root_), shoot (K_shoot_) and stem (K_stem_) hydraulic conductance were directly measured using a Hydraulic Conductance Flow Meter (HCFM; Tyree et al., 1993) (Dynamax Inc., Houston, TX, USA). Hydraulic parameters were determined in six loblolly pine and five bald cypress seedlings per treatment, and conductance values for a given plant were obtained from the same plant. To minimize the potential impact of diurnal periodicity on hydraulic conductance, all measurements were taken between 1000 hrs and 1200 hrs and under the same environmental conditions (temperature of 22 °C, and relative humidity of 60%). During HCFM measurements, the leaves were submerged in water to maintain constant temperature and prevent transpiration. To measure K_root_ and K_shoot_ the plants were cut 5 cm above the soil surface and the cut ends of the shoots and roots were connected to the HCFM. This instrument perfuses degassed water through root or shoot system by applying pressure to a water-filled bladder contained within the unit. K_root_ was determined between 2 and 4 minutes after shoot decapitation, thus minimizing measurement errors to less than 10% (Vandeleur et al. 2014; Rodriguez-Gamir et al., 2019; see also Supplementary Figure 1). The flow rate of water through root or shoot was determined under transient mode (Yang and Tyree, 1994), which consists in measuring flow rate under increasing pressure applied by a nitrogen gas cylinder. Transients were also performed on shoots after removal of leaves to determine K_stem_. The applied pressure gradually rose from 0 to 450 kPa over the course of approximately 1 minute and the flow rate at each pressure value was logged every 2 seconds using the Dynamax software. Hydraulic conductance (K) was then calculated using the formula: K=Q_v_/P; where Q_v_ is the volumetric flow rate (kg s^−1^) and P is the applied pressure (MPa). Hydraulic conductance was standardized to values for 25 °C to account for the effects of temperature on water viscosity. Because the HCFM operates under high pressure, the measured K_root_ and K_shoot_ represent maximum values of conductances, that is in the absence of embolized conduits. At the end of the measurements, all-sided leaf area of the shoots was determined with an LI-3100 leaf area meter (Li-Cor, Inc., Lincoln, NE, USA), and conductance values were expressed on leaf specific area basis (Yang and Tyree, 1994). All plant biomass fractions were then harvested and dried at 70°C for 48 hours and weighed. Further, mass-specific root hydraulic conductance (K_root_-biomass) was calculated by normalizing K_root_ by root dry mass.

Root (two opposite lateral roots per seedling taken about 2.5 cm down from the root collar were sampled) and stem tracheids were visualized by perfusing the decapitated samples with 0.1% toluidine blue and imaged at 90-180x magnification using a digital camera mounted on a widefield zoom stereo microscope (ZM-4TW3-FOR-9M AmScope, USA). Tracheid diameter was measured along four 4 radials rows per sample using an image analysis software (Motic Images version 3.2, Motic Corporation, China). In addition, the presence or absence of aerenchyma was assessed on 4 lateral roots per sample, which included the two used for tracheid size determination (no aerenchyma was present in the stems).

Whole plant hydraulic resistance was calculated as in Domec *et al.* (2016)

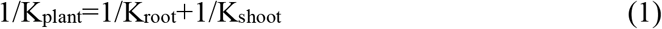

and resistances of the shoot components (1/K_stem_ and 1/K_leaf_) were calculated from the difference between resistances before and after removal of each leaf. Hydraulic conductance and resistance are reciprocals, and the latter is used for partitioning the resistances in the root-to-leaf continuum, and the former for examining the coordination between plant hydraulics and gas exchange.

### Aquaporin contribution to hydraulic conductances

We quantified the AQP contribution to K_root_ and K_shoot_ (and its components) and K_plant_ using hydroxyl radicals (*OH) produced using the Fenton reaction (solution made by equal mixing of 0.6 mM H_2_O_2_ and 3mM FeSO_4_) to inhibit AQP activity (Ye and Steudle, 2006; McElrone et al., 2007). Hydroxyl radicals has been shown to be less toxic and above all more effective in blocking water channels than mercuric chloride (Henzler and Steudle, 2004). Conductances with AQP function inhibited were measured by introducing approximately 18 ml of *OH solution, instead of water, into the existing compression couplings connected to the sample and the HCFM (McElrone et al. 2007; Johnson et al. 2014). As previously measured (Almeida-Rodriguez, Hacke and Laur 2011; Rodriguez-Gamir et al., 2019), the effect of *OH on conductivity was effective and reversable in less than 6 minutes when radicals were replaced with distilled water (Supplementary Fig. 1). Six transient curves per sample were constructed with the HCFM: two before inhibiting AQP activity, two after AQP inhibition, and two final ones after flushing the samples with water only to reassessed flow rate with no AQP inhibition. We calculated the AQP contribution to K_root_, K_shoot_ (K_stem_ and K_leaf_) and K_plant_ as the difference between initial conductance and conductance after AQP inhibition, divided by the initial conductance (Rodriguez-Gamir et al., 2019).

From measurements of conductances before and after inhibiting AQP activity, we were able to calculate whether the departure in values from control was due to either the xylem-only (structural changes in xylem conduits) or the AQP-only part of the hydraulic pathway. For a given stress applied, the structural part of the hydraulic pathway reducing conductance was calculated by dividing the difference in conductance between control and treatment after inhibiting AQP activity by the difference in conductance between control and treatment without inhibiting AQP activity. The AQP effect was taken as 1 minus the structural effect.

### Gas exchange and water potential

Net photosynthesis (A) and stomatal conductance (g_s_) were measured with a Li-Cor 6400 (Li-Cor, Inc., Lincoln, NE, USA). For each leaf, the chamber was set to match prevailing environmental conditions assessed immediately prior to the measurement: atmospheric CO_2_ concentration (390-410 ppm), relative humidity (46-59 %), photosynthetically active radiation (PAR; 1600-1800 μmol m^−2^ s^−1^), and leaf temperature (21-26 °C). All gas exchange results were expressed on an all-sided leaf area basis, and only fully-expanded healthy-appearing needles of the same age were picked for analysis. Maximum (light saturated) photosynthetic capacity (A_sat_) was measured on 4 green branchlets needles per seedling grown in the upper third of the plants. Immediately after the gas exchange measurements were performed, leaf water potential (Ψ_leaf_) was measured using a pressure chamber (PMS Ins., Albany, OR, USA). To assess maximum (least negative) Ψ_leaf_, two branchlets per individual from each treatment were sampled at predawn (between 05:00 hrs and 06:00 hrs).

Net photosynthesis versus intercellular CO_2_ concentrations (A-Ci curves) were measured at 25 °C leaf temperature, 60±10 % relative humidity and 1600 μmol m^−2^ s^−1^ PAR. The chamber CO_2_ concentrations were set to ambient and sequentially lowered to 50 ppm and then to 1500 ppm. These data were used to estimate the maximum Rubisco carboxylation (V_cmax25_), the maximum electron transport (J_max25_), and the dark respiration (R_d25_) rates according to Farquhar et al. (1980).

### Field hydraulic and canopy conductance

Two contrasting sites were used to determine field values of K_plant_ and g_s_ under typical field conditions, droughted and flooded conditions of large trees growing in intact forests. Soil salinity never occurred at the field sites to our knowledge. The first study site is a forested wetland located at the Alligator River National Wildlife Refuge, on the Albemarle–Pamlico Peninsula of North Carolina, USA (35°47’N, 75°54’W). This research site was established in November 2008, and includes a 35-m instrumented tower for eddy covariance flux measurements, a micrometeorological station, and 13 vegetation plots spread over a 4km^2^ area (Miao et al. 2013; Domec et al., 2015). The forest type is mixed hardwood swamp forest (>100-year-old); the overstory is predominantly composed of water tupelo (*Nyssa aquatica* L.) that represents 39% of the basal area and an even mix of red maple (*Acer rubrum* L.), bald cypress and loblolly pine. The canopy of this site is fairly uniform with heights ranging from 16 m to 21 m, and with leaf area index peaking at 4.0 ± 0.3 in early July.

The second, drier site (35°11’N, 76°11’W) located within the lower coastal plain, mixed forest province of North Carolina (Domec et al., 2009). This 100-ha mid-rotation (23-year-old) loblolly pine stand (US-NC2 in the Ameriflux database) was established in 1992 and has an understory comprised of other woody species such as sparse red maple and bald cypress trees. Artificial drainage lowers the height of the water table, improving site access and increases productivity, especially during winter months (Domec et al., 2015).

At both sites, canopy conductance was derived from sapflow measurements and thus comprises the total water vapor transfer conductance from the ‘average’ stomata of the canopy. Sapflow was measured at breast height using thermal dissipation probes inserted in two flood-adapted species (bald cypress and water tupelo) and two others not adapted to flooding (red maple and loblolly pine) (see Domec et al., 2015 for further description of the sites and the methodology used). Note that water tupelo was only present at the wetland site. Stomatal conductance of the plants measured in the field was calculated from transpiration and vapor pressure deficit (VPD), using the simplification of the inversion of Penman–Monteith model (Ward et al., 2013). To analyze the effect of K_plant_ on g_s_, K_plant_ from field and greenhouse samples was calculated from the slope of the relationship between diurnal variation in Ψ_leaf_ and transpiration (Loustau et al., 1995). Changes in Ψ_leaf_ from dawn to mid-afternoon were quantified with a pressure chamber (PMS, Albany, OR) on six to eight leaves collected from each tree equipped with sapflow sensors Oren et al. (1999) showed that under saturated light, the decrease in g_s_ with increasing VPD is proportional to g_s_ at low VPD. Therefore, the sensitivity of the stomatal response to VPD when PAR was above 800 μmol m^−2^ s^−1^ (light-saturated g_s_) was determined by fitting the data to the functional form:

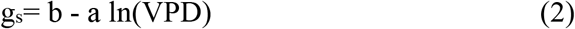

where b is g_s_ at VPD = 1 kPa (hereafter designated as reference or maximum canopy-averaged stomatal conductance, g_s-max_), and a is the rate of stomatal closure and reflects the sensitivity of g_s_ to VPD [dg_s_/dlnVPD, in mmol m^−2^ s^−1^ ln(kPa)^−1^]. We propose to use this framework where VPD and light intensity are fixed to investigate the nature of the relationship between K_plant_ and g_s-max_, and how this relationship affects the sensitivity of g_s_ to VPD.

### Statistical analyses

All measured parameters were tested using multiple analysis of variance with species, treatments, and AQP activity taken as factors. Mean separation was performed using the Tukey's procedure at 95 % confidence level. Statistical analyses were run using SAS (Version 9.4, Cary, NC, USA) and curve fits using SigmaPlot (version 12.5, SPSS Inc. San Rafael, CA, USA).

## Results

### Plant biomass

All treatments significantly reduced loblolly pine (*Pinus taeda* L.) total biomass (*p*<0.01; Table 1), whereas for the flood-tolerant bald cypress (*Taxodium distichum* L.) only the drought and the flooded plus salinity treatments had a negative effect on growth (*p*<0.037). This decrease in plant growth was mainly attributed to a reduction in root and stem biomass in bald cypress (*p*<0.032), and in leaf and stem biomass in loblolly pine (*p*<0.01). Despite this reduction in plant size, the fine root to leaf mass ratio of bald cypress was only affected in the flooding plus salinity treatment, whereas in loblolly pine it was stimulated by 25% and 55% in the flooding and the flooding plus salinity conditions, respectively. All stresses decreased leaf mass per area (LMA) in loblolly pine (*p*<0.02). In bald cypress LMA was only negatively affected by the drought and by the flooded plus salinity treatments (*p*<0.01). Field measurements indicated that unlike loblolly pine and red maple (*Acer rubrum* L.), flooded bald cypress and water tupelo (*Nyssa aquatica* L.) grew as rapidly (*p*>0.65) as trees subjected to periodic or non-flooded conditions (Supplementary Fig. 2).

**Table 1.** Plant, root (fine and coarse), stem and leaf dry masses (g), as well as mean tracheid diameters (D_t_stem_; D_t_root_), fine-root to leaf mass ratio (Root__fine_/Leaf), and leaf mass per area (LMA) for the different treatments of *Taxodium distichum* (n=5, +SE) *and Pinus taeda* (n=6, +SE). The presence (Yes - and the percentage of roots affected) or absence (No) of root aerenchyma (intercellular air spaces) observed 5 weeks after initiating the treatments is also indicated.

|  | Control |  | Drought |  | Flooded |  | Flooded plus Salt |  |
| --- | --- | --- | --- | --- | --- | --- | --- | --- |
|  | *T. distichum* | *P. taeda* | *T. distichum* | *P. taeda* | *T. distichum* | *P. taeda* | *T. distichum* | *P. taeda* |
| Plant (g) | 13.9 ± 1.1C | 10.7 ± 0.4B | 9.9 ± 0.9AB | 9.1 ± 0.4A | 11.7 ± 1.0BC | 9.5 ± 0.6A | 9.7 ± 0.8AB | 7.9 ± 0.8A |
| Root (g) | 6.1 ± 0.4B | 4.7 ± 0.4A | 4.4 ± 0.5A | 4.2 ± 0.3A | 5.0 ± 0.6AB | 4.9 ± 0.4A | 3.9 ± 0.5A | 4.3 ± 0.3A |
| Stem (g) | 5.4 ± 0.5D | 2.9 ± 0.2B | 3.3 ± 0.3C | 2.2 ± 0.1A | 3.9 ± 0.4 C | 1.9 ± 0.2 A | 2.8 ± 0.2B | 1.8 ± 0.1A |
| Leaf (g) | 2.4 ± 0.6AB | 3.1 ± 0.4B | 2.2 ± 0.4B | 2.6 ± 0.3B | 2.8 ± 0.8AB | 2.7 ± 0.3B | 2.9 ± 0.7AB | 1.7 ± 0.3A |
| Dt_stem (µm) | 23.1±3.8C | 14.8±2.1B | 19.9±2.2C | 9.4±1.2A | 22.6±4.1C | 13.6±1.5AB | 17.8±2.2BC | 10.2±1.5A |
| Dt_root (µm) | 12.3±0.9C | 10.2±1.0BC | 11.8±1.3C | 8.8±0.8AB | 13.3±1.6C | 9.0±0.6B | 10.7±1.1BC | 7.6±0.5A |
| Aerenchyma | Yes – 38 % | No | No | No | Yes – 84 % | No | Yes – 24 % | No |
| Root_fine/Leaf | 0.72 ± 0.09B | 0.53 ± 0.05A | 0.74 ± 0.10B | 0.50 ± 0.06A | 0.71 ± 0.12AB | 0.67 ± 0.04B | 0.48 ± 0.07A | 0.79 ± 0.9B |
| LMA (g cm<sup>-2</sup>) | 9.1 ± 0.9B | 13.4 ± 0.7D | 7.0 ± 0.6A | 11.8 ± 0.5C | 8.9 ± 1.0B | 11.4 ± 0.8C | 7.2 ± 0.4A | 9.0 ± 0.6B |

Values in horizontal sequences not followed by the same letter are significantly different at the 0.05 level.

### Effect of flooding and salinity on the partitioning of hydraulic conductance

All treatments decreased whole-plant hydraulic conductance (K_plant_) in loblolly pine (*p*<0.05), whereas bald cypress was only affected by the drought and the flooding plus salinity treatment (Fig. 1). In both species the strongest decrease in root (K_root_) and shoot (K_shoot_) hydraulic conductances were measured for plants subjected to flooding plus salinity. It should be noted that flooding alone did not affect any of the conductances in bald cypress. When loblolly pines were flooded, even K_root_ on a root-mass basis (K_root_biomass_) dropped significantly (by 45%), whereas K_root_ or K_root_biomass_ of the flood-tolerant bald cypress did not (Table 2; Fig.1).

**Figure 1:**
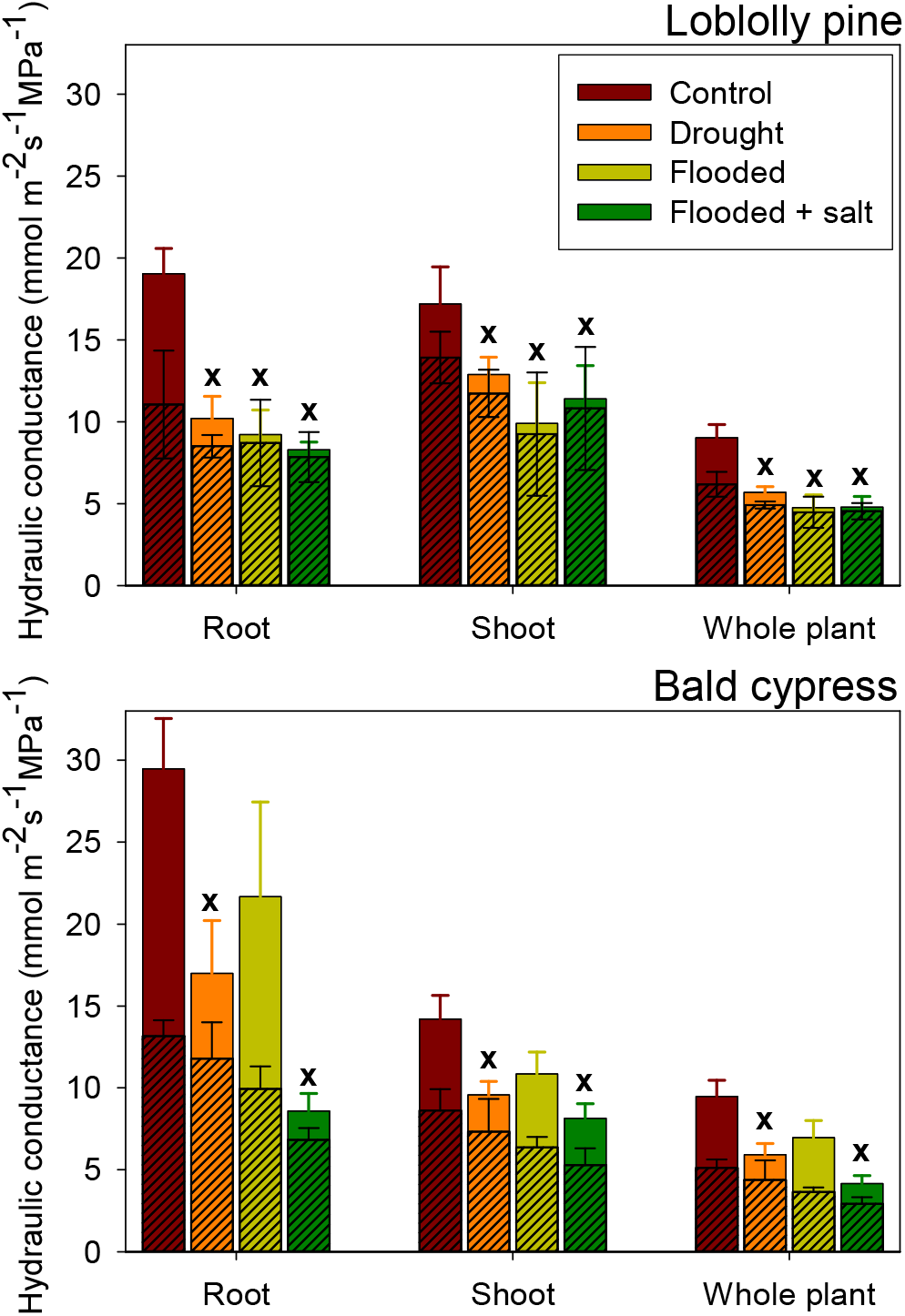
Mean values (+SE) of hydraulic conductances (solid bars) in root, shoot and whole loblolly pine (n=6) and bald cypress (n=5) plants growing in control, droughted, flooded and flooded + salt conditions. Crosses indicate a significant difference between control and any of the treatments (*p*<0.05). Hashed bars represent values of hydraulic conductance following aquaporin inhibition i.e. the xylem-only part of the hydraulic pathway.

**Table 2:**
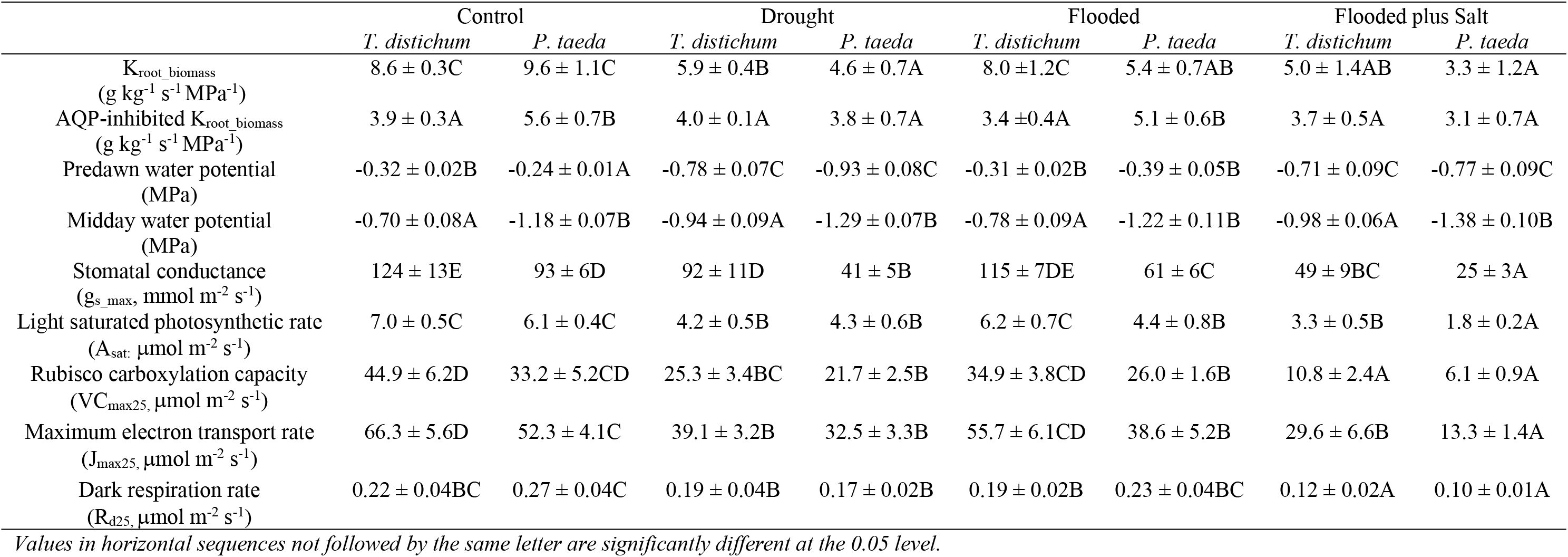
Mean root hydraulic conductance on a root-mass basis (K_root_biomass_), root hydraulic conductance on a root-mass basis after inhibiting aquaporin (AQP) activity (AQP-inhibited K_root_biomass_), leaf water potentials (Ψ_leaf_), stomatal conductance, light saturated photosynthesis, and photosynthetic parameters at 25°C (VC_max25_, J_max25_, R_d25_) for the different treatments of *Taxodium distichum and Pinus taeda*. Values are means +SE (n=5-6).

The overall decline in K_plant_ was mainly driven by an increase in root and stem resistances in loblolly pine, and by root resistance only in bald cypress (Fig. 2). Under control conditions, roots represented between 35% and 45% of whole-plant resistance (1/K_plant_), and under treatments this partitioning increased to more than 50% (*p*=0.038) in loblolly pine, which was paralleled by a reduction in leaf resistance from 35% to 15% (*p*=0.023). In bald cypress, only the flooded plus salt treatment increased the predominance of root resistance, which was accompanied by a decrease in the contribution of leaf and stem to the overall whole-plant resistance.

**Figure 2:**
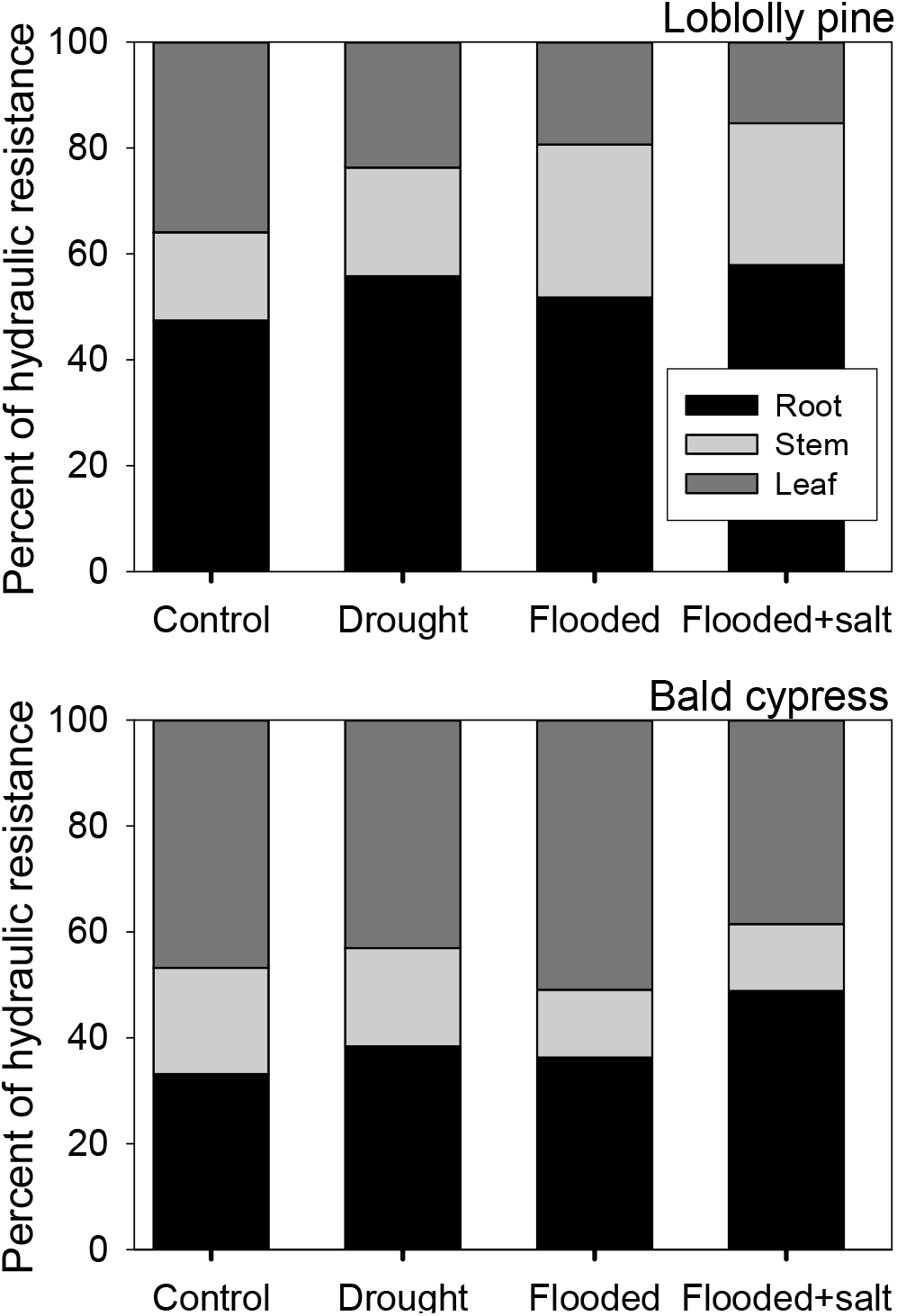
Partitioning of hydraulic resistances (1/conductance) of loblolly pine and bald cypress organs in control, flooded and flooded + salt conditions. Note that in all cases root and leaves represented more than 70% of total whole plant resistance.

### Aquaporin contribution to plant organ conductances and gas exchanges

The reduction in K_root_ and K_plant_ between control and the other treatments (Fig. 2) was mainly caused by a reduction in AQP activity rather than by a change in root anatomy (Table 1; Fig. 3). Even when K_root_ was calculated on a root-biomass basis (K_root-biomass_), the inhibition of AQP activity led to similar values of K_root-biomass_ (AQP-inhibited K_root_biomass_ in Table 2) across all treatments (*p*>0.47) in bald cypress, and for the flooded treatment (*p*=0.87) in loblolly pine. Nonetheless, in loblolly pine seedlings, AQP-inhibited K_root_biomass_ decreased by 31% (*p*=0.042) and 45 % (*p*=0.028) in the drought and flooded plus salt treatments, respectively, but that was still less than the overall reduction in K_root_biomass_ (52 % and 66 %, respectively), indicating that changes in K_root_biomass_ were mostly driven by the inhibition of AQP. This reduction in K_root_biomass_ in loblolly pine mirrored the decrease in root and stem tracheid diameter in the drought and flooded plus salt treatments (Table 1). In bald cypress, tracheid size was not affected by treatment, but aerenchyma production was stimulated under flooded conditions (Table 1).

**Figure 3:**
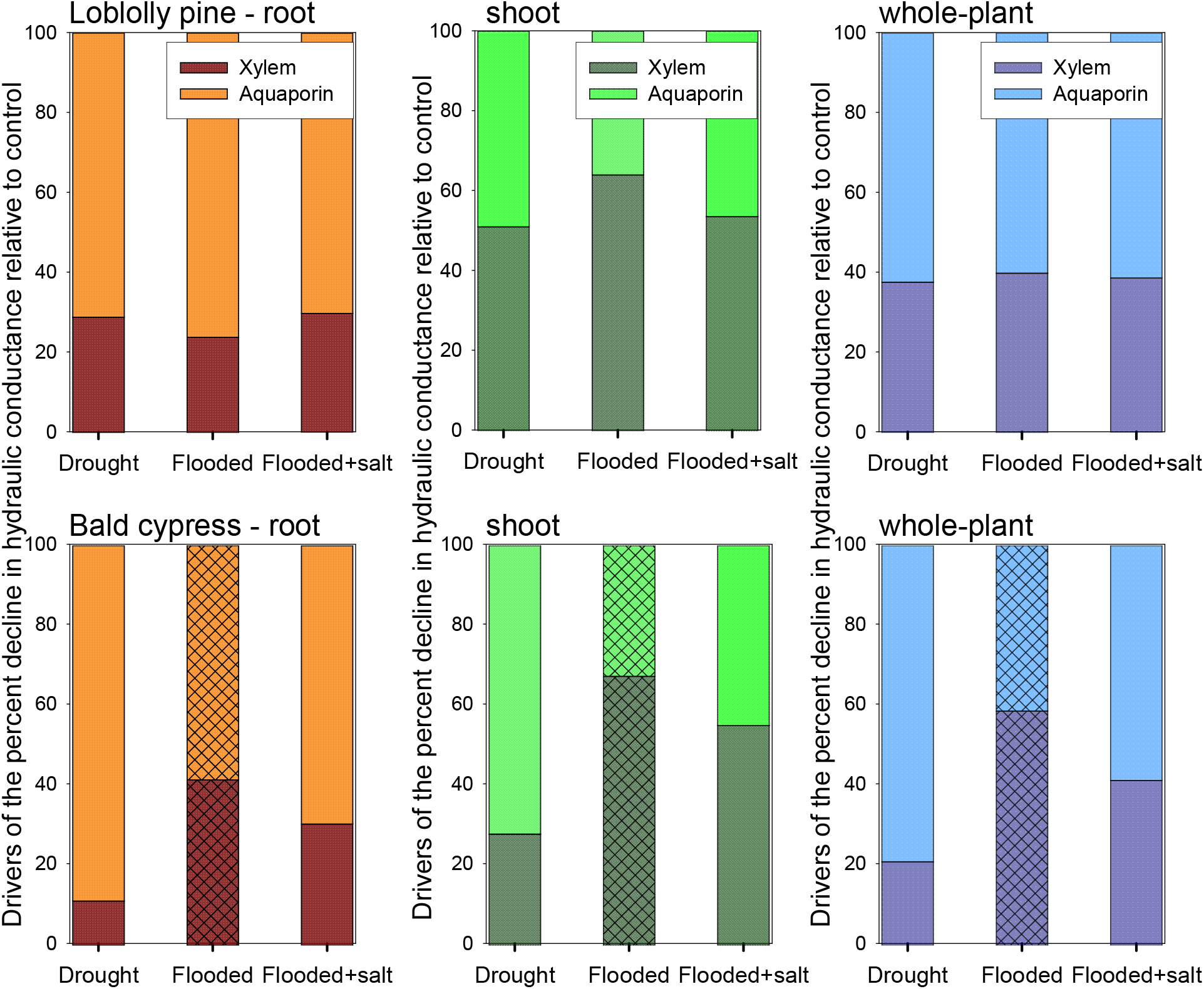
Effect (shown in percent) of either the xylem-only (structural changes in xylem conduits) or the aquaporin-only (AQP) part of the hydraulic pathway on the decrease in loblolly pine and bald cypress hydraulic conductance between control and drought, flooded, and flooded plus salinity treatments (absolute values of conductances are seen in Fig. 1). For a given stress applied, the structural part of the hydraulic pathway reducing conductance was calculated by dividing the difference in conductance between control and treatment after inhibiting AQP activity by the difference in conductance between control and treatment without inhibiting AQP activity. The AQP effect was taken as 1 minus the structural effect. Bars with patterns represent treatments that did not induce significant difference in conductance (*p*>0.05; flooded condition for bald cypress).

While blockage of AQP reduced hydraulic conductance, the extent of the decrease varied among organs and species (Fig. 3). Root AQP activity in loblolly pine decreased (*p*<0.001) from 42 % under controlled conditions, to less than 5-17 % in the different treatments, which was the driver of the decline in whole-plant AQP contribution (Fig. 4A). In this species, we found that flooding and flooding plus salinity reduced the AQP activity of the whole plants from 32 % to less than 6 % (*p*<0.01). In bald cypress only the drought and flooded plus salt treatments reduced (*p*<0.02) AQP contribution to K_root_ or K_plant_ (Fig. 4B). For that species, the inhibition of AQP in the flooding treatment did not affect (*p*=0.95) K_root_ or K_leaf_. In both species, drought also had a significant effect on AQP contribution to K_leaf_ with a decrease from 17 % to 9 % (*p*<0.03) in loblolly pine, and from 44 % to 23 % (*p*<0.001) in bald cypress. In both species there was no treatment effect on the contribution of AQP activity to K_stem_ (*p>*0.42).

**Figure 4:**
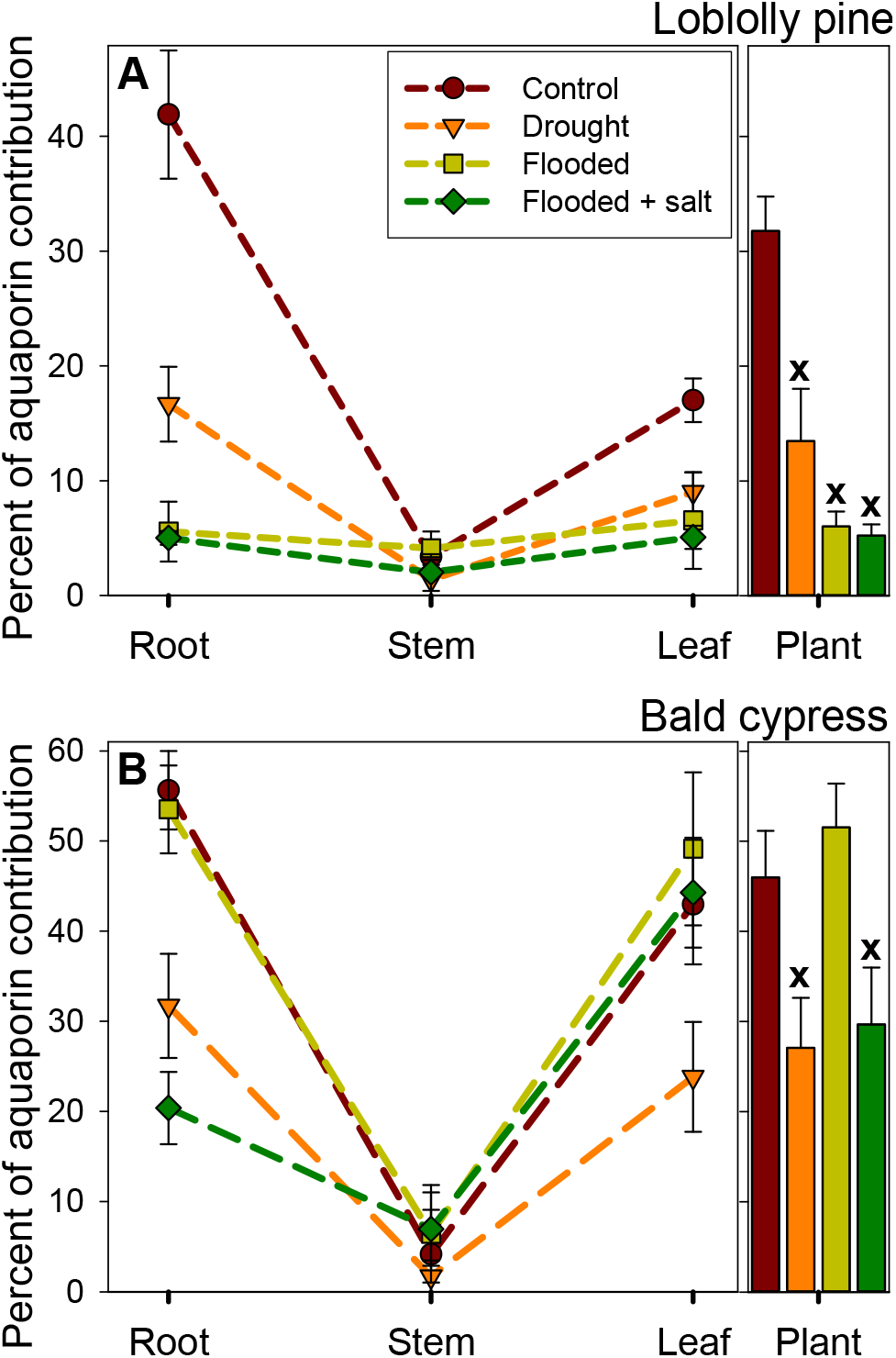
Aquaporin (AQP) contribution to root, stem, leaf and whole-plant hydraulic conductances in (A) loblolly pine (n=6, +SE) and (B) bald cypress seedlings (n=5, +SE) growing under control, water-stressed, flooded, and flooded plus salinity conditions. Crosses indicate a significant difference in whole-plant AQP contribution between control and any of the treatments (*p*<0.05).

Maximum stomatal conductance (g_s-max_; i.e*. g*s measured at a reference VPD of 1kPa and under saturated light) for bald cypress was only negatively affected by drought and the flooding plus salt treatment (Table 2). In loblolly pines, g_s-max_ differed under flooded and flooded plus salt treatments, experiencing the smallest and the largest stomatal closure, respectively. Photosynthetic rate of both species was also negatively affected by treatments (*p*<0.04), with the strongest reduction for the drought and flooded-salt treatments (Table 2). The disruption of photosynthesis concurred with a reduction in rubisco carboxylating enzyme activities and maximum electron transport rate (VC_max25_ and J_max25_, respectively; Table 2). Similarly, across species dark respiration rates were only affected by the flooded plus salt treatments.

After taking into account the effect of VPD on g_s_, K_plant_ had a major influence on g_s-max_ at field conditions. Across species and treatments, and whether plants were from the greenhouse or grown in the field, a 50% reduction in K_plant_ was accompanied by a 37 % decline in g_s-max_ (Fig. 5A). There was indeed no difference (*p*=0.33) in the relationship between g_s-max_ and K_plant_ for seedlings growing in greenhouse and mature trees in the field. Species differences were apparent in K_plant_, with higher values in bald cypress and red maple. Flooded loblolly pine exhibited the same level of reduced K_plant_ as water-stressed plants. In red maple, permanently flooded conditions reduced water uptake capacity more than two-fold, and this species exhibited higher hydraulic limitation and g_s-max_ in flooded than in drought-stressed conditions. However, bald cypress and water tupelo (circles and diamonds in Fig. 5A, respectively), which are species found in permanently wet soils, did not experience more than 15 % decline in g_s-max_ under flooded conditions. The sensitivity of g_s_ to VPD was linearly related to g_s-max_ (Fig. 5B) and K_plant_ (Supplementary Fig. 3). Stomatal conductance declined in response to increasing VPD, and the magnitude of the reduction varied over the measurement period as shown by the decline in g_s-max_. The slope of the relationship between g_s-max_ and the sensitivity of g_s_ to VPD (0.62 ± 0.04) was not different (*p*>0.99) than the previously reported generic value of 0.60 based on a hydraulic model that assumes tight stomatal regulation of Ψ_leaf_ (Oren et al., 1999).

**Figure 5:**
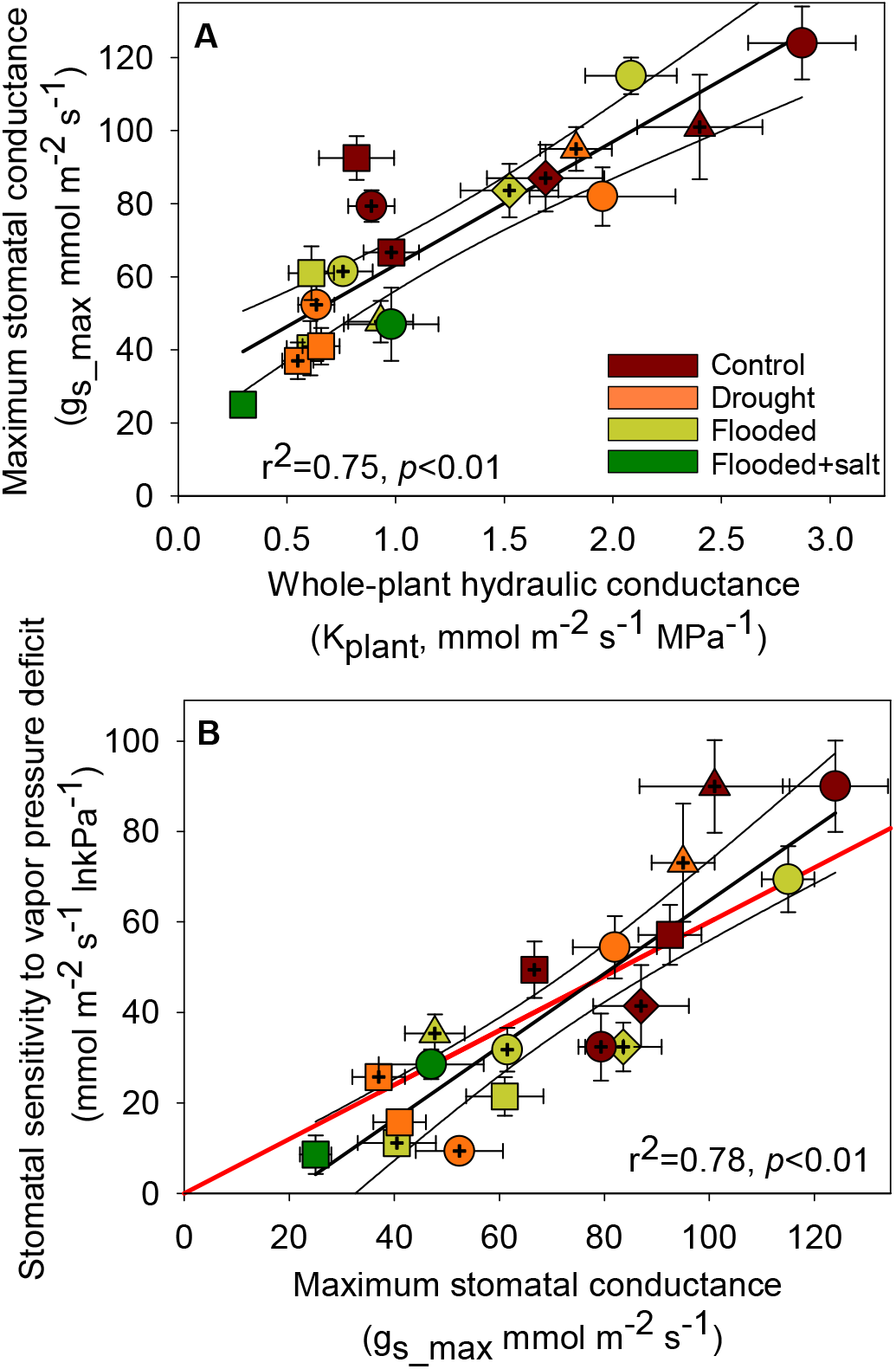
(A) Linear relationship between the maximum (reference) stomatal conductance (*g*_s_ at vapor pressure deficit = 1 kPa) and (B) plant hydraulic conductance (K_plant_) and between the sensitivity of stomatal conductance to vapor pressure deficit (dg_s_/dlnVPD) and g_s-max_ of plant species growing under control, water-stressed, flooded, and flooded plus salinity conditions. Circles, diamonds, squares and triangles represent bald cypress, water tupelo, loblolly pine and red maple, respectively. Crossed-filled symbols represent mature plants growing in the field, non-crossed symbols represent bald cypress and loblolly pine seedlings from the greenhouse experiment. In (B), the red line (slope = 0.6) indicates the theoretical slope between stomatal conductance at VPD = 1 kPa and stomatal sensitivity to VPD that is consistent with the role of stomata in regulating minimum leaf water potential (Oren et al. 1999).

Maximum g_s_ and stomatal sensitivity to VPD decreased linearly with increasing the contribution of root hydraulic resistance (1/K_root_) to 1/K_plant_ for both species (Fig. 6). Those negative relationships appeared also to be identical across treatments with a 50 % increase in resistance belowground resulting in a 56 % reduction in g_s_max_ and in a 65 % decrease in stomatal sensitivity to VPD.

**Figure 6:**
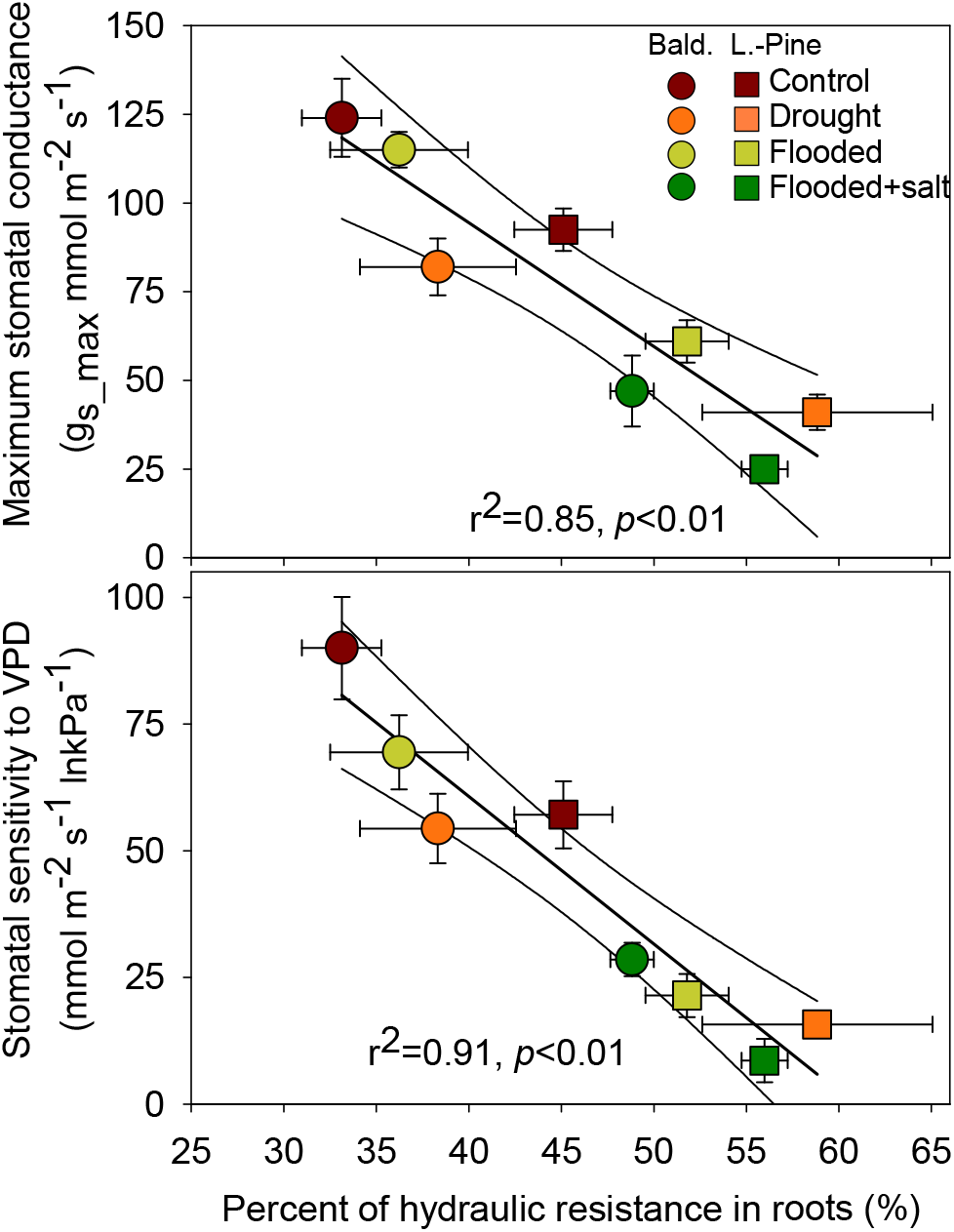
Maximum stomatal conductance (g_s-max_) and the sensitivity of stomatal conductance to vapor pressure deficit (dg_s_/dlnVPD) versus percent of hydraulic resistance in roots of bald cypress (Bald.) and loblolly pine (L.-Pine) seedlings growing under control, water-stressed, flooded, and flooded plus salinity conditions.

The decrease in g_s_max_ and A_sat_ were linked to a decrease in AQP contribution to root conductance among treatments and also species (*p*<0.039; Fig. 7). Although weaker, those relationships still held when whole-plant AQP activity was compared to gas exchange, and a 25 % decrease in AQP contribution to K_plant_ was predicted to reduce g_s_max_ by 38% and A_sat_ by 30 %.

**Figure 7:**
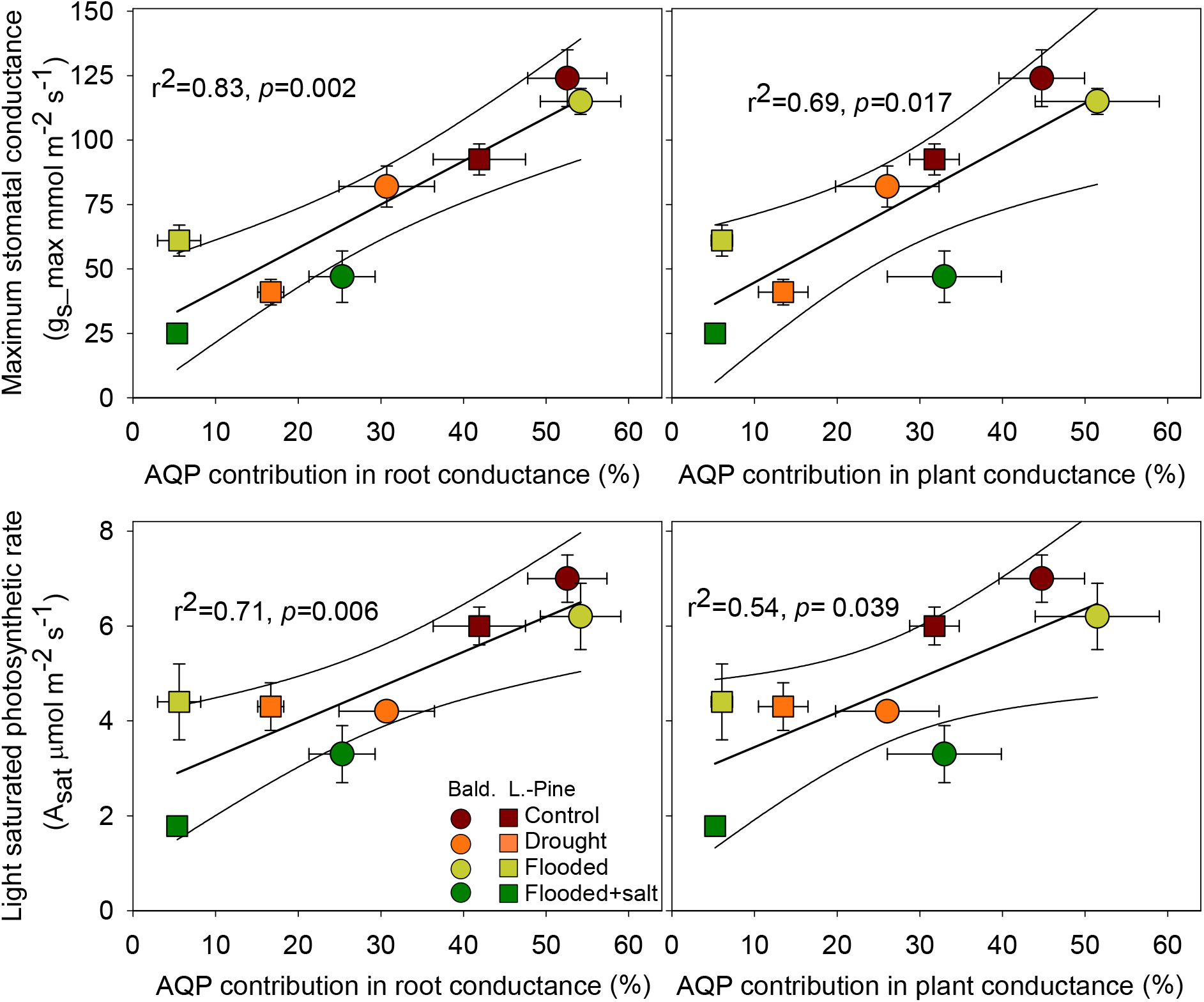
Maximum stomatal conductance (g_s-max_) and Light saturated photosynthetic rate (A_sat_) versus aquaporin (AQP) contribution to root, or whole-plant hydraulic conductances of bald cypress (Bald.) and loblolly pine (L.-Pine) seedlings growing under control, water-stressed, flooded, and flooded plus salinity conditions.

## Discussion

In US coastal regions from Maryland to Texas that are vulnerable to SLR (Titus and Richman, 2001; Kirwan and Gedan, 2019), many species such as bald cypress, water tupelo, red maple and loblolly pine are ecologically dominant and commercially important. The first two species are fully adapted to partial or total soil flooding and the other species are common to forest communities of estuarine woodlands (Pezeshki, 1992; Keeland and Sharitz, 1995). Bald cypress seedlings generally tolerate flooding, but marked with an initial reduction in growth (Allen et al., 1996). However, within 3 to 5 years, seedlings generally recover from the stress imposed by developing pneumatophores (Megonigal and Day, 1992), explaining why flooded trees may grow as rapidly as trees subjected to non-flooded conditions (Supplementary Fig. 2). However, before this root formation occurs, the influence of AQP on control of plant water movement and gas exchanges is needed and reflected in our results (Fig 4; Fig.7).

### Aquaporin activity appears to be essential in species-specific tolerance to stress

Our results highlight the integrated nature of hydraulics across the whole plant and emphasized the contributions of structural and physiological components to conductance (Bramley et al., 2009; Maurel and Nacry, 2020). Drought, flooding and flooding plus salinity treatments caused a significant shift in hydraulic resistance away from stem and leaves to the roots, because of differential transmembrane AQP activity and not because of changes in the apoplastic hydraulic pathway (xylem diameter) (Table 1, Fig. 3 and see AQP-inhibited K_root_biomass_ in Table 2). During stress, some structural and anatomical changes also occurred as seen by the decrease in either leaf, stem or root biomass under drought or flooded plus salt treatments, affecting for the latter treatment root to leaf area ratio in both species (Table 1). However and unlike the role played by AQP, those structural changes provided only minute adjustments in xylem hydraulic conductance (conductance once AQP activity was inhibited) and did not explain the whole decrease in conductance and thus the physiological mechanisms controlling water transport through the root (Table 2; Fig. 3). Both loblolly pine and bald cypress were highly susceptible to the combined stress of flooding plus salinity which lends support to the role of saltwater intrusion in the formation of coastal ghost forests, since bald cypress is also dying in these forests (Kirwan and Gedan, 2019). Lower predawn Ψ_leaf_ were expected with higher salinity because the lower osmotic potential of the medium (4 ppm corresponding to an osmotic potential of 0.31 MPa) likely increased leaf tissue ionic concentration (Allen et al., 1996). This excess of ions disrupted photosynthesis and inhibited carboxylating enzyme activities (Table 2), which in turn contributed to inhibited root or leaf growth, and the production of new aerenchymatous roots in the flood-adapted species (Table 1).

These findings may shed light on the adaptive advantages of altering AQP activity in response to environmental stresses (Maurel and Nacry, 2020). Regarding drought, lowering AQP activity in roots should lead to larger Ψ_leaf_ gradients, inducing stomata to close more rapidly. Reducing water channel activity can then be seen as a means to reduce water loss when soil water availability is low (McLean et al., 2011). In the case of flooding, the resulting decrease in K_root_ observed in the flood-intolerant species (such as loblolly pine used here) could also limit water transport to the leaves, causing stomatal closure and thus protecting the integrity of the whole hydraulic system until non-stressed conditions resume (Else et al., 2001). Loblolly pine is known to be tolerant to low salinity and short-term flooding (Poulter et al., 2008), and our experiment showed that this species reduced significantly gas exchange under these conditions, but to levels that were not lethal (Table 2). The negative impact of flooding on plants is a consequence of the low solubility of oxygen in water (Leyton and Rousseau, 1958), leading to anoxia (Kozlowski, 1997). The tight coupling of AQP functioning to the drop in cell energy (due to oxygen deprivation and acidosis) suggests that short-term adjustments in tissue hydraulics are critically needed during the early stages of the anoxic stress to balance water uptake with water loss (Tan et al., 2019). Long-term metabolic adaptation to flooding is generally characterized by the decrease in belowground biomass to limit oxygen deficiency, but one of the most adaptive features of plants of wetland ecosystems is aerenchymatous tissues characterized by intercellular gas-filled spaces that improve the storage and diffusion of oxygen. Unlike the adjustment in root biomass or xylem anatomy that can take more than 2 months (Krauss et al., 1999) and was not observed in any of the two seedlings (Table 1), intercellular air spaces, which were present after 5 weeks of flooding in bald cypress, likely played a vital role in maintaining root uptake and preventing the AQP-mediated reduction in K_root_. Kamaluddin and Zwiazek (2002) and Holbrook and Zwieniecki (2003) have also proposed that anoxia-induced AQP down-regulation may prevent the transport of ethylene precursors away from the root, thereby favoring the accumulation of ethylene to trigger the differentiation of root aerenchymas, especially in adapted species such as bald cypress (Table 1). Salinity added to the flooding stress may trigger larger AQP inhibition, so that advective salt flow to the root surface may be minimized (Azaizeh and Steudle, 1991). In the short term (a few hours following exposure to flooding or flooding plus salinity), it has been shown that plants respond to osmotic shock by reduced AQP activity (Martinez-Ballesta et al., 2000; Rodríguez-Gamir et al., 2012), which in our case was followed by reduced K_root_, most likely as an adaptive strategy to eliminate water loss from the roots under conditions of low osmotic potential.

Finally, it can also be hypothesized that the role of AQP may in fact not concern the primary response of the plant to stress, but its recovery performance (Siefritz et al., 2002). Stimulation of specific AQP suggested that “gating” in response to salt stress involved not only the reduction in water channels, but also an enhancement in the internalization of specific AQPs (raft–associated pathway), putatively becoming active once stress is relieved (Li et al., 2011).

### Root hydraulics as related to whole plant water transport and gas exchange

One of the objectives of our study was to evaluate a hypothesized correlation between leaf gas exchange and root hydraulics as influenced by AQP activity. The decline in g_s_max_ (and it is sensitivity to VPD) and photosynthesis was strongly related to the increase in root resistance due to a decrease in AQP contribution to K_root_, with a common relationship found among the species despite important differences in treatment responses (Figs. 6 and 7). In non woody plants, it has been suggested that abscisic acid (ABA) accumulation in leaves may be responsible for stomatal closure in flooded plants (Castonguay et al., 1993; Else et al., 2001). However, in woody plants the marked reduction in g_s_ in flooded seedlings does not seem to be induced by ABA, since a significant reduction in g_s_ appeared a week after stressors were applied (unpublished data; Rodríguez-Gamir et al., 2012), whereas the increase in ABA in leaves is generally detected 4-5 weeks later (Zhang and Zhang, 1994; Rodríguez-Gamir et al., 2012). Maximum g_s_ and K_plant_ were tightly coordinated in plants growing in the field or in greenhouse (Fig. 5). Changes in K_plant_, driven by K_root_, imposed a decline in g_s_max_, thus affecting leaf water status and further increases in transpirational water loss and carbon assimilation. Midday Ψ_leaf_ did not change during the flooding treatment, highlighting the adaptive role of stomatal closure in counteracting leaf dehydration (Meinzer, 2003). Furthermore, our data indicate that flooded pines exhibited the same level of reduced K_plant_ as water stressed plants. Field data showed that red maple exhibited higher hydraulic limitation and higher g_s-max_ in flooded than in water stressed conditions, indicating that species differences exist in the response to flooding. In contrast, bald cypress and water tupelo regulated very efficiently the closure of stomata, thus adjusting the evaporative water losses to the water uptake capacity of roots and the resulting decrease in K_plant_ (Fig. 5).

Our results also showed that the sensitivity of g_s_ to VPD was mostly attributable to the variation in g_s-max_, which is consistent with the isohydric regulation of Ψ_leaf_ induced by K_plant_ (Oren et al., 1999). Stomata responded to VPD in a manner consistent with protecting xylem integrity and thus the capacity for water transport (Domec et al., 2009; McCulloh et al., 2019). Future climate change is expected to increase temperature and therefore VPD in many regions (Oppenheimer et al., 2019). Stomatal acclimation to VPD as affected by drought, flooding and flooding plus salinity could potentially have a large impact on the global water and carbon cycles.

Here we measured that in forested wetlands global plant transpiration responses to future climate will probably not differ from expectations based on the well-known relationship between g_s_ and VPD (Oren et al., 1999). To improve climate predictions of warming effects on transpiration for plants subjected to different abiotic stresses, our results indicated that modelers could potentially allow for predictable shifts in g_s_ under water stress but also flooded conditions, combined with the use of single coefficient conveying g_s_ sensitivity to VPD.

Woody species responses to flooding and flooding plus salinity are wide-ranging and can change based on the life-history stage of a plant. Seedlings are generally more sensitive to salinity while mature plants may show a wider range of tolerance (Kozlowski, 1997). However, when our field and greenhouse observations were analyzed together, some common responses were observed (Fig. 5), highlighting the need for integrating data on seedlings and mature plants in future studies on wetland adaptation to SLR and its restoration (Carmichael and Smith, 2016).

### Conclusion

Our study provides new functional and mechanistic insights on plant hydraulics by showing that the components of K_plant_ are highly dynamic, reflecting a balance between species adaptive capacity and AQP functioning. Neither species tolerated flooding plus salinity. In loblolly pine, high water uptake was largely mediated by active transport through AQP, but was easily disrupted by drought, flooding and salinity. In bald cypress, a flooded-tolerant species, AQP contribution to water transport was less sensitive overall and did not respond to flooding. Under controlled conditions, AQP activity and xylem structure were colimiting root water transport. However, in response to environmental factors, except again for the flooding treatment in bald cypress, AQP functioning rather than changes in xylem structure or biomass allocation controlled the fluctuations in K_root_, and thus in K_plant_. The decline in K_leaf_ was rather the consequence of both a decrease in AQP activity and in structural changes. An important challenge was also to integrate the AQP-mediated reduction in K_root_ within the mutual interactions of roots and shoots and its putative effect on gas exchange. As such, across species and treatments, the reduction in *g*_s_ and its sensitivity to VPD appeared to be direct responses to decreased K_plant_ and was influenced by AQP contribution to water transport.

## Data availability statement

The data supporting the findings of this study are available from the corresponding author (Jean-Christophe Domec) upon request.

## Author contributions

J.-C.D. and D.M.J. conceived the original screening and research plans; J.-C.D., D.M.J., R.W. and M.J.C. performed the hydraulics experiments, A.T.O., M.J.C. and R.W. performed the gas exchange experiments; J.S.K., J.-C.D., A.N., and G.M. performed field experiments; W.K.S. provided plant materials; J-C.D. and D.M.J. analyzed the data and wrote the article with contributions of all the authors.

